# scExplorer: A Comprehensive Web Server for Single-Cell RNA Sequencing Data Analysis

**DOI:** 10.1101/2024.11.11.622710

**Authors:** Sergio Herńandez-Galaz, Andrés Hernández-Oliveras, Felipe Villanelo, Alvaro Lladser, Alberto J.M. Martin

## Abstract

Single-cell RNA sequencing (scRNA-seq) has redefined our ability to investigate cellular heterogeneity at unprecedented resolution. However, the computational analysis required for scRNA-seq data interpretation remains a significant barrier for many researchers. We present scExplorer, an interactive web application that streamlines the analysis workflow from preprocessing through dimensional reduction to differential expression analysis. The platform provides an intuitive interface for real-time data exploration and visualization while maintaining compatibility with established analysis frameworks in both R and Python environments. scExplorer significantly reduces the technical knowledge and computational requirements needed to perform scRNA-seq analysis, enabling researchers to focus on biological insights rather than computational complexity. The application modular design ensures scalability and robust performance, making it a valuable tool for both small-scale and large-scale single-cell studies.

**Graphical Abstract:** 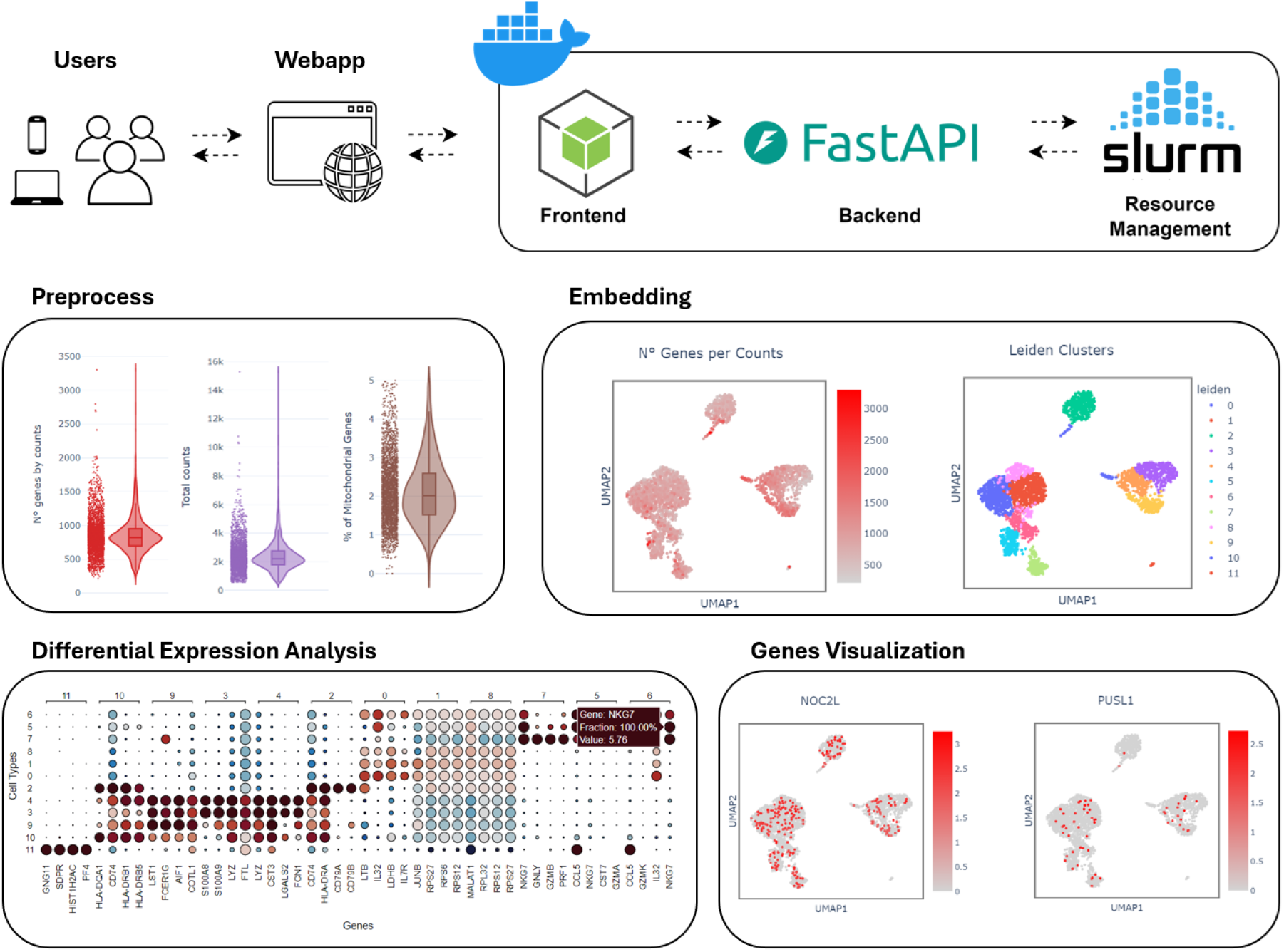

## 1 Introduction

Advances in next-generation sequencing technologies have enabled the development of single-cell RNA sequencing (scRNA-seq), a powerful approach that captures gene expression profiles at individual cell resolution. Since the introduction of this technique in 2009, scRNA-seq has facilitated the identification of novel cellular markers and provided insights into continuous processes critical to developmental biology and various physiological phenomena [1, 2, 3]. The field has experienced remarkable growth, as evidenced by the exponential increase in publications indexed in PubMed, expanding from under 100 papers in 2009 to about 5,000 publications by the end of 2024. This dramatic surge in scientific output reflects the technology increasing adoption and importance across biological disciplines. In parallel to this publication trend, there has been substantial growth in publicly available scRNA-seq datasets. Data repositories such as the Gene Expression Omnibus (GEO) [4] and the Human Cell Atlas Data Portal [5] have seen a significant expansion in their data collections. For instance, the Single Cell Expression Atlas [6] hosts thousands of experiments comprising millions of cells across diverse species and tissue types, making it an invaluable resource for the scientific community that is expected to continue its steady growth in the future. Building on the increasing relevance of scRNA-seq, several tools have been developed to analyze scRNA-seq data systematically. Prominent among these are Seurat in R [8] and Scanpy in Python [7].However, the requirement for programming skills remains a major obstacle, preventing many scientists with strong biological expertise and relevant research questions from effectively interpreting their data. scExplorer was developed specifically to overcome this barrier, simplifying the analysis process and enabling researchers to focus on biological insights rather than computational challenges.

## 2 Materials and Methods

The scExplorer web application offers a complete pipeline for scRNA-seq analysis, addressing key challenges associated with data quality, dimensionality reduction, clustering, and differential expression. Figure 1. It supports data import in multiple formats, including .h5ad, .h5, .rds, and 10x Cell Ranger outputs, allowing for easy interconnection with most used data generation and processing pipelines.

**Figure 1:**
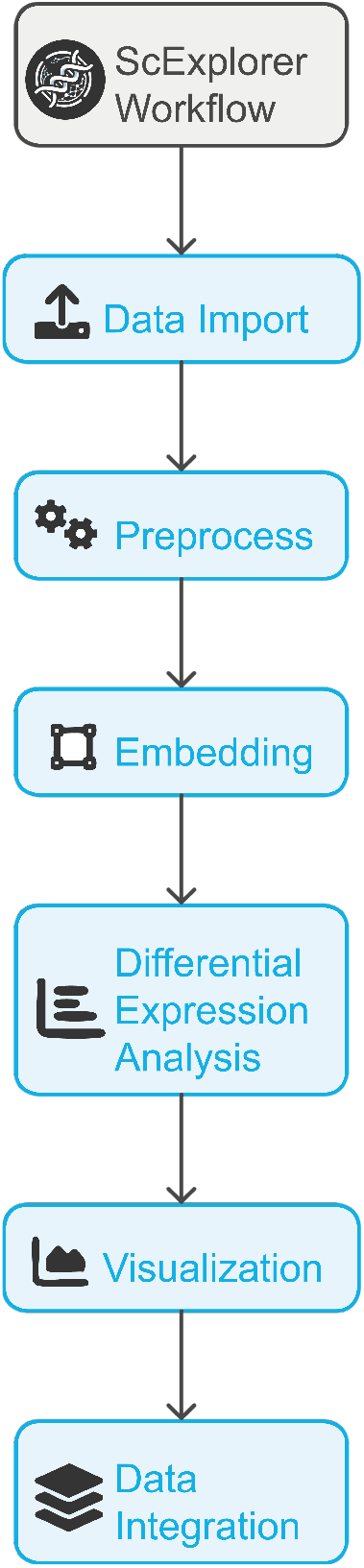
Overview of scExplorer Features. scExplorer streamlines scRNA-seq analysis with Scanpy (Python) and Seurat (R). It performs data import, quality control, normalization, and selection of high variable genes. PCA and UMAP visualize cell relationships, Leiden clustering identifies groups, and differential expression analysis finds genes markers for each cell type identified. Combat, Scanorama, BBKNN, and Harmony correct batch effects for data integration.

scExplorer incorporates standard quality control procedures, ensuring in this way that only high-quality data proceeds to analysis by filtering low-quality cells and non-informative genes, reducing technical noise and improving the reliability of downstream results. During preprocessing, cells with insufficient gene expression are removed to retain biologically relevant data, while filtering out genes expressed in very few cells avoids statistical artifacts. Cells with excessive mitochondrial gene expression, often indicative of cellular stress, are excluded by default to maintain data integrity. Doublet detection using Scrublet prevents merged cells from being misinterpreted as unique populations, preserving the resolution of single-cell data [9].

Identification of the most significant sources of variation, i.e. highly variable genes (HVGs), ensures that the analysis captures the most informative features. In scExplorer, users can select between the Seurat and Cell Ranger methods for HVG identification. The method implemented in Seurat stabilizes variance across datasets, particularly in multi-batch analyses, while the approach available from Cell Ranger ensures optimal performance for datasets generated using 10x Genomics pipelines.

UMAP embedding is used to visualize high-dimensional data in two-dimensional space, revealing both local and global cellular patterns [10]. The size of the local neighborhood influences the structure captured by UMAP, while usage of PCA components improve computational efficiency. scExplorer utilizes the Leiden algorithm for clustering, an approach that optimizes modularity to identify communities of cells based on their similarity. [11]. The resolution parameter controls the granularity of clusters, providing flexibility to explore both broad populations and fine-grained subpopulations. To assist with optimizing clustering results, scExplorer integrates Clustree [12], a tool for visualizing cluster resolutions across different parameter values. Clustree helps users systematically assess how varying the resolution impacts the number of cell types, enabling more informed choices about the appropriate resolution for their specific dataset. With this approach, scExplorer helps define biologically distinct populations in the dataset.

Differential expression analysis (DEA) is performed to identify key marker genes for each cluster of cells analyzed. scExplorer offers for this purpose the Wilcoxon rank-sum test and the t-test to accommodate different data distributions. The Wilcoxon test, being non-parametric, is effective for scRNA-seq data with non-normal distributions, while the t-test offers higher power for normally distributed data. DEA results can automatically highlight the top-ranking genes for each cluster or evaluate a user-defined list of genes, making scExplorer support both exploratory and targeted investigations.

All intermediate results, including dimensionality reductions, cluster annotations, and differential expression findings, are exportable in Python (.h5ad) and R (.rds) formats, facilitating further analysis. Interactive visualizations built with Plotly enhance data exploration and presentation [13].

scExplorer modular architecture, based on Docker containers, isolates components such as the frontend (Node.js/Express), backend (FastAPI), analysis modules (Scanpy and Seurat), and job scheduling (SLURM). This design ensures scalability, simplifies maintenance, and enables efficient resource management, supporting reliable performance for both small and large datasets.

### 2.1 Batch correction for multidataset integration

Batch effects, often introduced by differences in sample preparation, sequencing protocols, or technical artifacts, represent a significant challenge in scRNA-seq data analysis. These effects can mask true biological variation and lead to spurious clustering or incorrect downstream interpretations. Correcting for batch effects is essential to ensure that datasets from different sources can be integrated seamlessly, enabling robust identification of shared and unique cell populations across conditions or experiments.

To address this challenge, scExplorer incorporates multiple state-of-the-art tools for batch correction,such as Combat [14], Scanorama [15], batch-corrected k-nearest neighbors (BKNN) [16] and Harmony [17], each tailored to specific needs:

#### Combat

Originally designed for microarray data, Combat uses parametric and non-parametric empirical Bayes framework for adjusting batch effects in scRNA-seq data. This method is integrated into scExplorer to allow users to harmonize datasets from different sources, ensuring that downstream analyses such as clustering and trajectory inference are reflective of biological rather than technical variations.

#### Scanorama

this tool corrects batch effects by aligning disparate datasets using mutual nearest neighbors, which are identified through a dimensionality-reduced space. It is particularly useful for large-scale integrations where datasets exhibit significant heterogeneity.

#### BKNN

this method mitigates batch effects by adjusting the graph-based relationships in the data. It uses local neighborhood information from each batch to perform a more refined integration, enhancing the detection of subtle biological variations across different conditions.

#### Harmony

this algorithm operates by adjusting the low-dimensional embeddings of the cells. It iteratively corrects for batch effects while preserving the biological signal, making it a powerful tool for datasets with complex batch structures.

Each of these batch correction methods can be selected and customized in the scExplorer web interface, allowing users to tailor their analysis according to the specific challenges and characteristics of their datasets.

### 2.2 Pre-uploaded datasets and Tutorial

scExplorer comes pre-loaded with three example datasets to demonstrate its functionality: Human PBMC, a standard benchmark dataset for scRNA-seq workflows; Mouse cortex, hippocampus, and subventricular zone cells, useful for exploring brain-specific cellular diversity; and Zebrafish cranial neural crest-derived cells, ideal for studying developmental biology and lineage tracing [18]. For testing and validation, we showcase the analysis of human PBMCs through the entire workflow, as illustrated in Figure 2.

**Figure 2:**
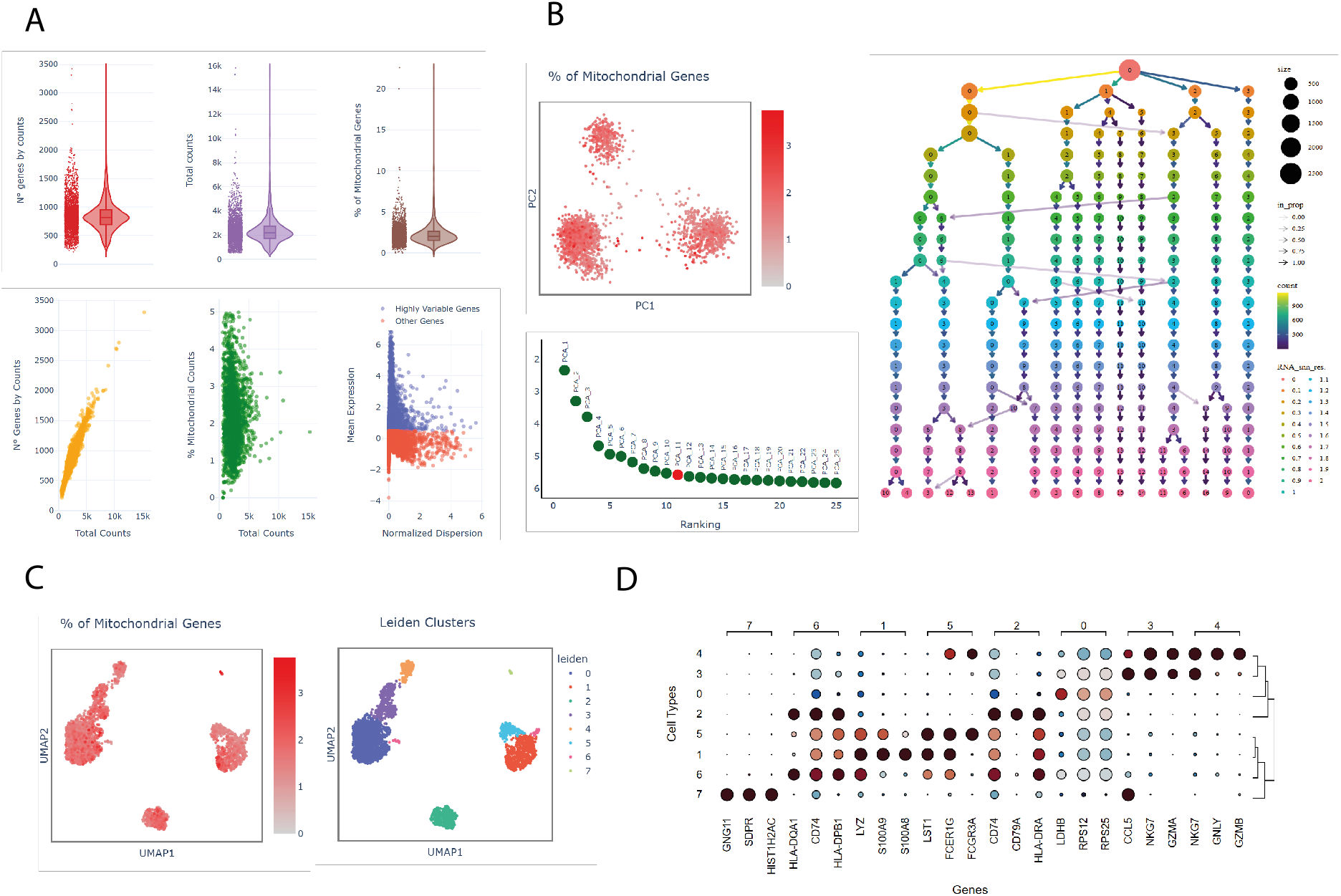
scExplorer pipeline. A) Summary of quality control metrics for scRNA-seq data, which assessed key aspects of individual cell quality to ensure reliable downstream analysis. The top row displayed violin plots summarizing the number of detected genes per cell (left), total UMI (Unique Molecular Identifier) counts per cell (middle), and the percentage of mitochondrial gene counts (right). The first scatter plot in the bottom row related the number of detected genes with total UMI counts. The second scatter plot examined the relationship between the percentage of mitochondrial gene expression and total counts, and the final scatter plot identified highly variable genes against normalized dispersion. B) Principal Component Analysis (PCA) plot showed cells colored by the percentage of mitochondrial genes. Below it, an elbow plot ranked the principal components by their variance contribution, helping to identify the most informative components for Uniform Manifold Approximation and Projection (UMAP). On the right, a hierarchical clustering graph visualized cell clusters for optimal cell type identification. C) UMAP plot showed cells colored by the percentage of mitochondrial genes. On the right, clusters were identified by the Leiden community detection algorithm. D) dot plot presented the results of differential expression analysis for the identified clusters on the dataset, showing the log fold change of key genes.

Additionally, scExplorer features an in-depth tutorial designed to guide users through the interface and explain each parameter in detail. This tutorial not only facilitates efficient use of the tool but also provides educational resources for understanding scRNA-seq analysis principles, empowering researchers with both technical and theoretical knowledge.

## 3 Discussion

The development and deployment of scExplorer addresses critical challenges in single-cell RNA sequencing (scRNA-seq) data analysis. While scRNA-seq technology has advanced rapidly, the computational expertise required for data analysis often creates a bottleneck in research workflows. Several platforms exist for single-cell RNA-seq (scRNA-seq) analysis, including CellxGene, ASAP, and Loupe Cell Browser, each offering different strengths and trade-offs. CellxGene, developed by the Chan Zuckerberg Initiative, excels in real-time visualization of large-scale single-cell datasets through UMAP and t-SNE embeddings, providing intuitive exploration tools without requiring programming expertise. However, it is primarily designed for data exploration and lacks a comprehensive computational pipeline for tasks like preprocessing or batch correction, which are essential for raw scRNA-seq data analysis.

Similarly, ASAP offers web-based tools for interactive scRNA-seq analysis but focuses more on the annotation and comparison of known datasets rather than enabling users to perform detailed preprocessing on new datasets. Loupe Cell Browser, part of the 10x Genomics ecosystem, provides in-depth visualization of 10x Genomics data but is limited to users working within that specific framework.

In contrast, scExplorer offers an end-to-end solution, integrating Python and R frameworks to support preprocessing, quality control, dimensional reduction, clustering, and differential expression analysis. Unlike the other platforms, scExplorer enables users to conduct doublet detection with Scrublet, batch correction using several tools and flexible export options in python and R formats. Additionally, its microservices-based backend ensures scalability, facilitating the processing of both small and large datasets through SLURM job scheduling. This makes scExplorer a powerful alternative that bridges the gap between user-friendly interfaces and robust computational capabilities, helping researchers perform comprehensive analyses without extensive coding knowledge.

## 4 Acknowledgements

The author gratefully acknowledges Juan Manuel Cabello for his valuable assistance with the network communication aspects and his support in the deployment of scExplorer.

## 5 Code Availability

scExplorer is already accessible at http://apps.cienciavida.org/scexplorer, allowing users to perform single-cell RNA sequencing (scRNA-seq) data analysis directly through a web interface without the need for local installations.

For users interested in deploying their own instance or contributing to the project, the source code is open and freely available on GitHub at https://github.com/networkbiolab/scexplorer.

